# Environmental DNA analysis shows high potential as a tool for estimating intraspecific genetic diversity in a wild fish population

**DOI:** 10.1101/829770

**Authors:** Satsuki Tsuji, Atsushi Maruyama, Masaki Miya, Masayuki Ushio, Hirotoshi Sato, Toshifumi Minamoto, Hiroki Yamanaka

**Affiliations:** Graduate School of Science, Kyoto University, Kitashirakawa-Oiwakecho, Sakyo-ku, Kyoto 606–8502, Japan; Faculty of Science and Technology/Graduate School of Science and Technology, Ryukoku University, 1-5 Yokotani, Seta Oe-cho, Otsu, Shiga 520–2194, Japan; Department Ecology and Environmental Sciences, Natural History Museum and Institute, Chiba, Chiba 260–8682, Japan; Hakubi Center, Kyoto University, Yoshida-honmachi, Sakyo-ku, Kyoto, 606-8501, Japan; Center for Ecological Research, Kyoto University, 2–509–3 Hirano, Otsu 520–2113, Japan; PRESTO, Japan Science and Technology Agency, Kawaguchi 332–0012, Japan; Graduate School of Human and Environmental Studies, Kyoto University, Sakyo, Kyoto, 606–8316, Japan; Graduate School of Human Development and Environment, Kobe University, 3-11 Tsurukabuto, Nada-ku, Kobe 657–8501, Japan; Center for Biodiversity Science, Ryukoku University, 1–5 Yokotani, Seta Oe-cho, Otsu, Shiga 520–2194, Japan

**Keywords:** Environmental DNA, intraspecific genetic diversity, mitochondrial haplotype, wild fish population, Field water

## Abstract

Environmental DNA (eDNA) analysis has recently been used as a new tool for estimating intraspecific diversity. However, whether known haplotypes contained in a sample can be detected correctly using eDNA-based methods has been examined only by an aquarium experiment. Here, we tested whether the haplotypes of Ayu fish (*Plecoglossus altivelis altivelis*) detected in a capture survey could also be detected from an eDNA sample derived from the field that contained various haplotypes with low concentrations and foreign substances. A water sample and Ayu specimens collected from a river on the same day were analysed by eDNA analysis and Sanger sequencing, respectively. The 10 L water sample was divided into 20 filters for each of which 15 PCR replications were performed. After high-throughput sequencing, denoising was performed using two of the most widely used denoising packages, UNOISE3 and DADA2. Of the 42 haplotypes obtained from the Sanger sequencing of 96 specimens, 38 (UNOISE3) and 41 (DADA2) haplotypes were detected by eDNA analysis. When DADA2 was used, except for one haplotype, haplotypes owned by at least two specimens were detected from all the filter replications. This study showed that the eDNA analysis for evaluating intraspecific genetic diversity provides comparable results for large-scale capture-based conventional methods, suggesting that it could become a more efficient survey method for investigating intraspecific genetic diversity in the field.

## Introduction

The field of environmental DNA (eDNA) analysis has rapidly developed over the past decade and has become a useful approach for investigating the species distribution of aquatic macroorganisms (e.g. Ficetola et al., 2008; Jerde et al., 2011; Takahara et al., 2013; Rees et al., 2015; Tsuji et al., 2019). Environmental DNA analysis enables us to detect the presence of species efficiently and non-invasively by analysing eDNA contained in water samples instead of capturing or observing the target species (Rees et al., 2014; Fukumoto et al., 2015; Thomsen et al., 2015; Yamanaka & Minamoto 2016). Combined with high-throughput sequencing (HTS) technology, eDNA metabarcoding is recently being used to provide information on species diversity by massively determining DNA sequences in eDNA samples (e.g. Miya et al., 2015; Shaw et al., 2016; Thomsen et al., 2016; Ushio et al., 2017; Yamamoto et al., 2017). More recently, researchers have begun to apply eDNA analysis to estimate intraspecific genetic diversity in a population (Sigsgaard et al., 2016, Parsons et al., 2018).

Nonetheless, two major issues remain to be solved before the practical application of eDNA-based methods for estimating intraspecific genetic diversity in wild populations: (1) the overestimation of intraspecific genetic diversity due to numerous sequences derived from PCR amplification and sequencing errors (cf. Schloss et al., 2011; Coissac et al., 2012; Edgar et al., 2016), and (2) the lack of information on the usefulness (e.g. detection efficiency [Can we evaluate intraspecific genetic diversity with less labour than capture-based methods?] and accuracy [Can we detect haplotypes corresponding with captured specimens?]) under field conditions. It is difficult to distinguish erroneous sequences from the original sequences because of their close similarity and hidden true genetic diversity. Sigsgaard et al. (2016) focused on the detection of only known haplotypes and tried to eliminate error sequences by clustering sequences into operational taxonomic units (OTUs, OTU methods). However, this strategy will fail in using the potential of eDNA analysis that allows for the detection of massive haplotypes including the unknown ones. Furthermore, in previous studies (Sigsgaard et al., 2016, Parsons et al., 2018), water sampling for eDNA analysis and tissue collection for Sanger sequencing were independently performed over a long period, and the survey period was different between the two surveys. Thus, when water sampling was performed, it is unclear whether there were individuals with haplotypes consistent with those detected by eDNA analysis.

To address the first issue, Tsuji et al. (2019) proposed an analytical strategy that combined the denoising of HTS data and subsequent haplotype selection on the basis of detection probability among multiple replications. This study aimed to accurately detect all known haplotypes contained in tank water. For the denoising of HTS data, two popular denoising software packages based on different denoising strategies, UNOISE3 (Edgar et al., 2016) and Divisive Amplicon Denoising Algorithm 2 (DADA2, Callahan et al., 2016), were used to examine the effect of the difference of denoising strategy. To denoise HTS data, each denoising software package emphasises on the sequence abundance and number of differences between sequences or the quality scores of the reads, respectively. As a result, regardless of the denoising software package used, the proposed denoising approach successfully eliminated > 99% of false-positive haplotypes derived from PCR and sequencing errors. However, further field studies are still needed to make a robust conclusion on the usefulness of the proposed analytical strategy for evaluating intraspecific genetic diversity because only tank water was used in Tsuji et al. (2019).

Regarding the second issue, to apply eDNA analysis for the evaluation of intraspecific genetic diversity on routine biodiversity surveys, we need to examine how efficiently the eDNA-based methods can evaluate intraspecific genetic diversity in the population at the sampling site. Tsuji et al. (2019) reported that we could detect all multiple known haplotypes derived from eDNA of tank water; however, field water usually contains more varying haplotypes with low concentrations, PCR inhibitors, and foreign substances. Thus, there is still uncertainty about the usefulness of the eDNA-based method for evaluating intraspecific genetic diversity from field water. The examination of the usefulness of eDNA-based methods for evaluating intraspecific genetic diversity under field conditions will aid in the further development of this method and survey design decisions. Molecular analyses to obtain more detailed insights into population genetic diversity at the individual level (e.g. Restriction-site Associated DNA Sequencing [RAD-seq], Multiplexed ISSR Genotyping by sequencing [MIG-seq], and Genome-Wide Association Study [GWAS]) cannot be applied to eDNA samples because eDNA is usually a mixture of multiple individuals ’ DNAs. However, these molecular analyses require tissue samples and the labour for capture survey. Thus, when eDNA-based methods provide comparable results for large-scale of the capture-based conventional method by only water sampling, we can find great value in the use of the eDNA-based methods for evaluating intraspecific genetic diversity.

This study aimed to examine whether we could accurately and effectively evaluate intraspecific genetic diversity in wild fish populations using an eDNA-based method by comparing the results with those of the capture-based conventional method. We performed water sampling and capture survey at a river on the same day and evaluated the intraspecific genetic diversity on the basis of eDNA analysis and Sanger sequencing of tissue DNA, respectively. In eDNA analysis, to examine the relationship between the detection probability and the frequency of each haplotype in the specimens, we prepared multiple replications for filtration and PCR. Also, we performed denoising using UNOISE3 and DADA2 and compared the results to examine a more suitable denoising method for eDNA-based evaluation of intraspecific genetic diversity. We explored the potential for detecting intraspecific genetic diversity with less labour and cost by comparing accumulation curves of detected haplotypes which were estimated using all or some data of PCR replications per filter.

## Materials and Methods

### Sampling sites and experimental design

In this study, Ayu (*Plecoglossus altivelis altivelis*), an important fishery target in Japanese inland waters (Iguchi et al. 2002, Takeshima et al. 2016), was used to examine the efficiency and accuracy of the eDNA-based method for evaluating intraspecific genetic diversity. A water sample and Ayu specimens were collected in the lower reach of Ado River (35°19’30" N, 136°03’49" E), a tributary of Lake Biwa, Japan, on 1 May 2015, when numerous individuals of Ayu were migrating upstream from Lake Biwa. The Ayu specimens were captured by a local fisherman within 12 h before water sampling using a fishing weir located approximately 30 m uphill from the water sampling site. We purchased 96 Ayu specimens from the local fisherman after water sampling. To avoid the cross-contamination of Ayu eDNA, different investigators collected water samples and purchased Ayu, and each water sample and Ayu specimens were stored separately. The entire experimental design is shown in Fig. 1. Per the current laws and guidelines of Japan relating to fish experiments, collection of fish tissue for DNA extraction and the use of DNA samples are allowed without the need for ethical approvals from authorities.

**Fig. 1.**
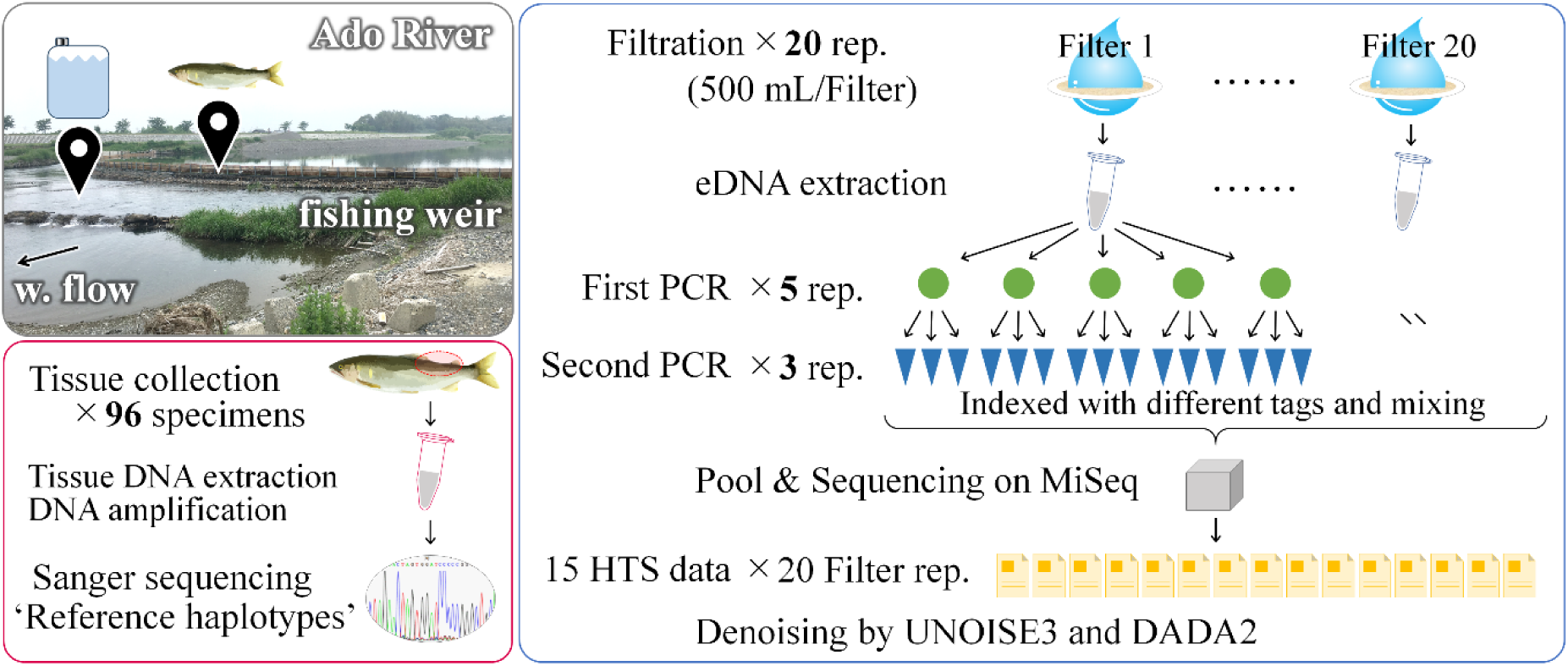
Experimental design.

### Determination of sequence from captured specimens by Sanger sequencing

A total of 96 specimens of Ayu (11.43 ± 1.84 g wet weight, mean ± SD) were preserved using ice immediately after capture. Approximately 0.02 g of the skeletal muscle tissue was excised from each specimen in the laboratory, and DNA was extracted from the tissue using DNeasy Blood & Tissue Kit (Qiagen, Hilden, Germany) following the manufacturer’s protocol. The tissue DNA was finally eluted in 200 µL of Buffer AE provided in the kit. For Sanger sequencing, almost the entire control region of mitochondrial DNA was amplified using PaaDlp-1_F (5’-GCTCCGGTTGCATATATGGACC-3’) and PaaDlp-1_R (5’-AGGTCCAGTTCAACCTTCAGACA-3’) (Tsuji et al., 2019) by StepOnePlus Real-Time PCR System. The 20-µL reaction mixture contained 900 nM of PaaDlp-1(F/R) in 1 × PCR master mix (TaqMan Gene Expression Master Mix), and 1 µL of tissue-derived DNA (10 ng/µL). The thermal cycle profile was 2 min at 50ºC, 10 min at 95ºC, 44 cycles of 15 s at 95ºC, and 60 s at 60ºC. Sequences of the PCR amplicons were determined by commercial Sanger-sequencing service (Takara Bio Inc. Kusatsu, Japan). Sequences of all reference haplotypes are listed in Table S1 and deposited to the DNA database of Japan (Accession number: LC433925–LC433966).

### Water sampling and filtration for eDNA analysis

Ten-liters of surface water was collected using a pristine plastic tank. The water quality parameters were measured using separately sampled water and water quality sensors (HI 98130; HANNA Instruments, Woonsocket, RI, USA). The water qualities were as follows: pH 6.35, temperature 17.4 °C, and electrical conductivity 0.05 mS/cm. Immediately after agitation of the whole 10 L water sample, 500 mL of sub-sample was filtered on-site onto each of 20 glass fibre filters (Whatman GF/F, 0.7-μm mesh, GE Healthcare, Chicago, USA). As a filtration negative control, the 500 mL of ultrapure water was filtered on-site and treated in the same manner in the following experiments as the real samples. The filter discs were immediately stored at −20 °C until eDNA extraction. To avoid contamination, all sampling and filtering equipment were soaked in 10% bleach solution for more than 10 min, carefully washed with tap water, and finally rinsed with ultrapure water before use.

### eDNA extraction from filter samples

Environmental DNA was extracted from the filter samples following the procedures of Tsuji et al. (2019). Each filter disc was placed into a spin column (EZ-10 SpinColumn & Collection Tube; Bio Basic Inc., Ontario, Canada), from which a silica-gel membrane was prospectively removed, and excess water on the filter was removed by centrifugation. Then, the mixture, containing 200-µL ultrapure water, 100-µL buffer AL, and 10-µL proteinase K, was added onto the filter and incubated at 56ºC for 30 min. After centrifuged at 6000 × *g* for 1 min. Then, 200 µL of TE buffer (pH 8.0) was added onto each filter to recover residual DNA on the filter and incubated at room temperature for 1 min. After centrifuged at 6000 × *g* for 1 min, 200 µL of Buffer AL and 600 µL of 100% ethanol were added. The mixture was then applied to a DNeasy Mini Spin Column, which was supplied by DNeasy Blood & Tissue Kit, and centrifuged at 6000 × *g* for 1 min. The DNA was purified according to the manufacturer’s instructions, and finally, the DNA was eluted with 100 µL of Buffer AE instead of the manufacturer’s recommended 200 μL. The extracted DNA samples were stored at −20 °C. The reagents, Buffer AL, Buffer AE and proteinase K, which were used for DNA extraction, were attachment reagents from DNeasy Blood & Tissue Kit.

### Library preparation and paired-end sequencing by MiSeq

Paired-end library preparation and MiSeq sequencing were performed using the same method as described by Tsuji et al. (2019). The two-step tailed PCR approach was employed to construct paired-end libraries. The first-round PCR (1st PCR) was performed using a primer pair that amplifies the control region of Ayu mitochondrial DNA species specifically. The primer pairs were developed in Tsuji et al. (2019) and contained adapter sequences and random hexamers (N). The sequence of primers are as follows: PaaDlp-2_F (5’-ACACTCTTTCCCTACACGACGCTCTTCCGATCTNNNNNNCCGGTTGCATATATGGACCTATTAC-3’), PaaDlp-2_R1 and R2 (5’-GTGACTGGAGTTCAGACGTGTGCTCTTCCGATCTNNNNNNGCTATTRTAGTCTGGTAACGCAAG −3’). The R indicates A (PaaDlp-2_R1) or G (PaaDlp-2_R2). The PCR mix with a total volume of 12 µL contained: 300 nM of PaaDlp-2_F, 150 nM each of PaaDlp-2_R1 and R2 in 1 × KAPA HiFi HotStart ReadyMix (KAPA Biosystems, Wilmington, MA, USA), and 3 µL sample eDNA. The 1st PCR was performed with five replicates for each eDNA sample derived from 20 filter replications. Additionally, triplicated PCR negative controls were included for each PCR run to monitor cross-contamination during the library preparation. The thermal cycle profile was 3 min at 95ºC, 35 cycles of 20 s at 98ºC, 15 s at 60ºC, and 15 s at 72ºC followed by the final extension for 5 min at 72ºC. The 1st PCR products were purified using the Agencourt AMPure XP (Beckman Coulter, USA) according to the manufacturer’s instructions (reaction rate: AMPure beads 0.8: PCR product 1, target amplicon length: ca. 290-bp).

The second-round PCR (2nd PCR) was performed in triplicates for each replication of the 1st PCR product (total 15 replications for each filter sample). The final PCR mix with a volume of 12 µL contained 300 nM of each 2nd PCR primer in 1 × KAPA HiFi HotStart ReadyMix, and 3 µL of the purified 1st PCR product. Samples were distinguished from each other on the bases of different combinations of indexing primers in the bioinformatics analysis. The primer sequences used in the 2nd PCR are listed in Table S2. For the PCR negative controls of the 2nd PCR, the PCR negative controls in the 1st PCR were added as a template. The thermal cycle profile was 3 min at 95ºC, 12 cycles of 20 s at 98ºC and 15 s at 65ºC, with the final extension for 5 min at 72ºC.

All indexed 2nd PCR products were pooled in equal volumes (1 µL each), and the target size of the libraries (ca. 370-bp) was collected using 2% E-Gel Size Select Agarose Gels (Thermo Fisher Scientific, USA) according to the manufacturer’s instructions. DNA concentrations of the collected libraries were adjusted to 4 nM (assuming 1 bp of DNA has the molecular weight of 660 g) using ultrapure water. Finally, the libraries were sequenced on a single MiSeq run using an Illumina MiSeq Reagent Kit v2 for 2 × 150 bp PE (Illumina, San Diego, CA, USA). All sequencing reads obtained in the present study were deposited in the DNA Data Bank of Japan (DDBJ) Sequence Read Archive (accession number: DRA009149).

### Bioinformatics analysis using denoising software packages

The full range of amplicon obtained using PaaDlp-2 primers (166 bp) was successfully sequenced by the MiSeq platform; however, after the forward primer of PaaDlp-1, some bases remained undetermined by the Sanger sequencing of the tissue-derived DNA. Scince the forward primers of PaaDlp-2 (for HTS) and PaaDlp-1 (for Sanger sequencing) had designed at close sites, three bases after the forward primer of PaaDlp-2 required omission to compare the overlapping regions between the two datasets. Thus, only 163 bp, which was successfully determined by both methods, was used for the subsequent bioinformatics analyses.

To perform denoising of erroneous sequences using software packages, the fastq files containing raw reads, which were obtained from 20 filters with 15 PCR replicates, were processed using the UNOISE3 (Edgar 2016) and DADA2 package ver. 1. 12. 1 (Callahan et al., 2016) of R. The UNOISE3 and DADA2 are two of the ASV methods for denoising sequencing errors in Illumina-sequenced amplicons, and 99.8% (UNOISE3) and 99.3% (DADA2) of false-positive haplotypes were successfully eliminated in a tank water experiment (Tsuji et al., 2019). The UNOISE3 and the DADA2 differ in how they correct sequencing errors. UNOISE3 uses a one-pass clustering strategy that does not depend on quality scores. The denoising algorithm of UNOISE3 considers sequence abundance and the number of differences between sequences to predict whether a sequence was correct or not (Edgar, 2016). The denoising algorithm of DADA2 corrects the sequence on the bases of the error model that is trained using quality scores of the reads and the probability of various copy errors that could be introduced during PCR amplification and sequencing. See Edgar (2016) and Callahan et al. (2016) for a full description of each ASV method. The UNOISE3 and DADA2 use alpha option (UNOISE3) and OMEGA_A and OMEGA_C (DADA2) as parameters to control the stringency of the denoising step. In the previous study using tank water (Tsuji et al., 2019), the default values (alpha = 2, OMEGA_A = 1e-40, and OMEGA_C = 1e-40) recovered the known haplotype composition of the sample better than did other values that we examined. However, it was suggested that further examinations of the effect of these parameters should be performed using field water samples. Thus, in this study, we preliminarily explored optimal values for alpha in UNOISE3 and OMEGA_A in DADA2, respectively. We used default values for OMEGA_C of DADA2 because we previously found that the result was mostly unaffected by these changes. In UNOISE3, we found that an alpha = 3 exhibited better performance than did the default value (alpha = 2) (see Discussion and Table S3). Conversely, in DADA2, we found that the default value of OMEGA_A showed better performance than when other values were used (see Table S2). Thus, in this study, UNOISE3 and DADA2 were performed with alpha = 3 and default settings with the other parameters, respectively.

### Denoising by UNOISE3

For UNOISE3, Usearch v 11 was used (Edgar 2010). Paired reads were merged using ‘fastq_mergepairs’ command and stripped of both the primer sequences and three base pairs from forward read. After quality filtering using the ‘fastq_filter’ command, fasta files containing all the unique sequences and their abundance were generated using the ‘fastx_uniques’ command. Finally, the detected unique sequences were denoised using the UNOISE3 (Edgar, 2016) option. The threshold of the minimum abundance and length filter were set at 4 reads and 100 bp, respectively. To identify chimaeras, the UCHIME2 (Edgar, 2016) option was used.

### Denoising by DADA2

For DADA2, fastq files containing raw reads were processed using the DADA2 package ver 1. 12. 1 (Callahan et al., 2016) and R ver. 3. 6. 0 software (R Core Team., 2019). The trimming of primer and random hexamers were performed using the function ‘removePrimers’. Reads with one or more expected errors (maxEE = 1) were discarded during a quality inspection, and three base pairs were trimmed from the forward read. The error model was trained using the function ‘learnErrors’ and used for sample inference of dereplicated reads using the function ‘dada’. Finally, paired reads were merged using the function ‘mergePairs’. In DADA2, the function ‘removeBimeraDenovo’ is implemented to identify chimeras. However, when it was used in this study, some haplotypes that were detected by the conventional method based on Sanger sequencing were incorrectly identified as chimeras. The algorithm of ‘removeBimeraDenovo’ might incorrectly identify haplotypes of Ayu included in the sampled water as chimaeras because of high sequence similarity. Thus, the identification and elimination of chimaera were not performed.

### Data analysis

All statistical analyses were performed using the statistical software R ver. 3. 6. 0 and the significance level were set at *p* ≤ 0.05. To examine the relationship between the number of analysed specimens and detected haplotypes in the conventional method (hereafter ‘reference haplotype’), the accumulation curve was estimated using the function ‘specaccum’ in the vegan v.2.5-5 package (Oksanen et al., 2018) of R. Additionally, sample size‐based rarefaction and extrapolation sampling curves were estimated using the function ‘iNEXT’ in the iNEXT v. 2.0.19 package (Hsieh et al. 2016) of R. Total sequence reads of each haplotype detected by UNOISE3 and DADA2 were compared using linear regressions.

The results obtained by DADA2 were used in the subsequent analysis because it indicated better detection accuracy than did the results of UNOISE3 (see Results). Generalised linear models (GLMs) with logit link function assuming binomial distribution were used to analyse the relationships between the detection probability of each haplotype among the 20 filters and 15 PCR replicates and the number of specimens that owned each haplotype in the conventional method (hereafter ‘owner specimens’). To examine the relationship between the number of detected reference haplotypes and the number of the analysed filters with different numbers of PCR replicates, the accumulation curves were estimated using two data sets, namely, (1) 15 PCR replications per filter (five 1st PCR replications × three 2nd PCR replications) and (2) five PCR replications per filter (five 1st PCR replications and only the first replication out of three 2nd PCR replications). Additionally, for reference haplotypes, the total sequence reads of each haplotype and the number of owner specimens were compared using linear regressions.

## Results

### Haplotypes detected by the conventional method

In the Sanger sequencing of tissue-derived DNA, a total of 42 reference haplotypes were detected from 96 Ayu specimens. The number of detected reference haplotypes increased with the number of analysed specimens, but the accumulation curve of detected reference haplotypes did not reach a plateau even after all 96 specimens were analysed (Fig. 2A). The number of owner specimens of each haplotype ranged from 1 (reference haplotype ID; 12–42) to 28 (reference haplotype ID: 01) (Fig. S1). Sample-based rarefaction and extrapolation sampling curves in a conventional method suggested that haplotype diversity at the sampling site was about 250 types (Fig. 2B).

**Fig. 2.**
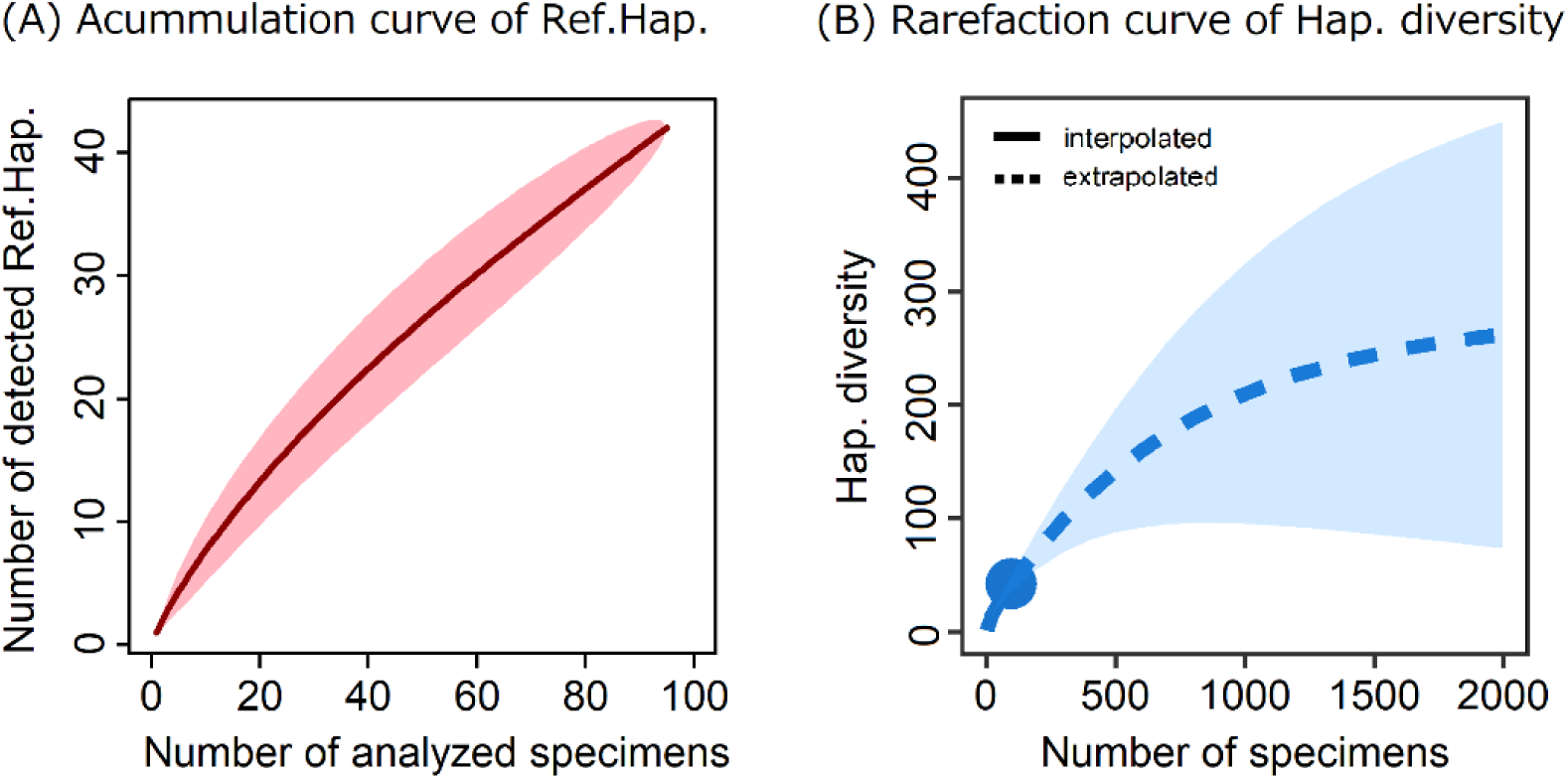
(A) Accumulation curve of reference haplotypes (the detected haplotypes from 96 specimens of Ayu by Sanger sequencing) based on the number of analyzed specimens and (B) sample-based rarefaction and extrapolation sampling curves in a conventional method. The solid and dashed lines in (B) indicate rarefaction and extrapolation, respectively. The shaded area corresponds to the 95% confidence intervals.

### Comparison of haplotypes detected by eDNA analysis using UNOISE3 and DADA2

In MiSeq paired-end sequencing of eDNA, a total of 20,377,762 reads were obtained from the 333 libraries (including 300 real samples, 15 filtration negative controls, and 18 PCR negative controls).

After denoising using UNOISE3 with alpha = 3, a total of 11,187,431 reads remained and were successfully assigned to 972 haplotypes. Of 42 reference haplotypes, 38 haplotypes were detected using the 20 filters (Fig. S1A). The multiple specimen-owned reference haplotypes were detected from 3–20 filters (average 17.3 filters) with a detection probability of 0.13–1 (average 0.8), whereas the single specimen-owned reference haplotypes (27 reference haplotypes) were detected from 1–20 filters (average 13.5 filters) with a detection probability of 0.09 to 1 (average 0.7), among the 15 PCR replications. The other unknown 934 haplotypes that did not correspond to the reference haplotypes were also detected from 1–20 filters (average 3.4 filters) with a detection probability of 0.07–1 (average 0.2) among the 15 PCR replications. The filtration negative controls and PCR negative controls yielded 942 reads, which were assigned to 25 haplotypes. All the sequence reads detected from negative controls were omitted in the subsequent analysis because of the negligible read abundance (<0.008% total reads).

After denoising using DADA2, 14,370,583 reads remained, and those were successfully assigned to 967 haplotypes. Of the 42 reference haplotypes, 41 haplotypes were detected from the 20 filters with 15 PCR replications (Fig. S1B). The 11 multiple specimen owned reference haplotypes were detected from all 20 filters except for one (reference haplotype ID 8, detected from one filter). The 30 single specimen-owned reference haplotypes were detected from 4–20 filters (average 18.1 filters) with a detection probability of 0.08–1 (average 0.7) among the 15 PCR replications. The other 926 reference non-corresponding haplotypes were detected from 1–20 filters (average 5.1 filters) with a detection probability of 0.07–1 (average 0.2) among the 15 PCR replications. In filtration negative controls and PCR negative controls, 2,398 reads were detected and finally assigned to five haplotypes. All sequence reads detected from negative controls were omitted in the following analysis because read abundance was negligible (<0.02% total reads).

Of the 42 reference haplotypes, 38 reference haplotypes were detected by both UNOISE3 and DADA2, 3 types were detected by DADA2 only, and 1 reference haplotype was undetectable by both methods. Each of the reference haplotypes showed a significant positive relationship in total read abundance between the two denoising software packages (lm, *p* < 0.001, R^2^ = 0.85, Estimate 0.34; Fig. 3A). A total of 1,428 haplotypes did not correspond to any of the reference haplotypes, and their total read abundance showed a significant positive relationship between the two denoising software packages (lm, *p* < 0.001, R^2^ = 0.10, Estimate 0.36; Fig. 3B). The 430 haplotypes shared by UNOISE3 and DADA2 also exhibited a significant positive relationship in total read abundance between the two denoising software packages (lm, *p* < 0.001, R^2^ = 0.85, Estimate 1.02; Fig. 3B).

**Fig. 3.**
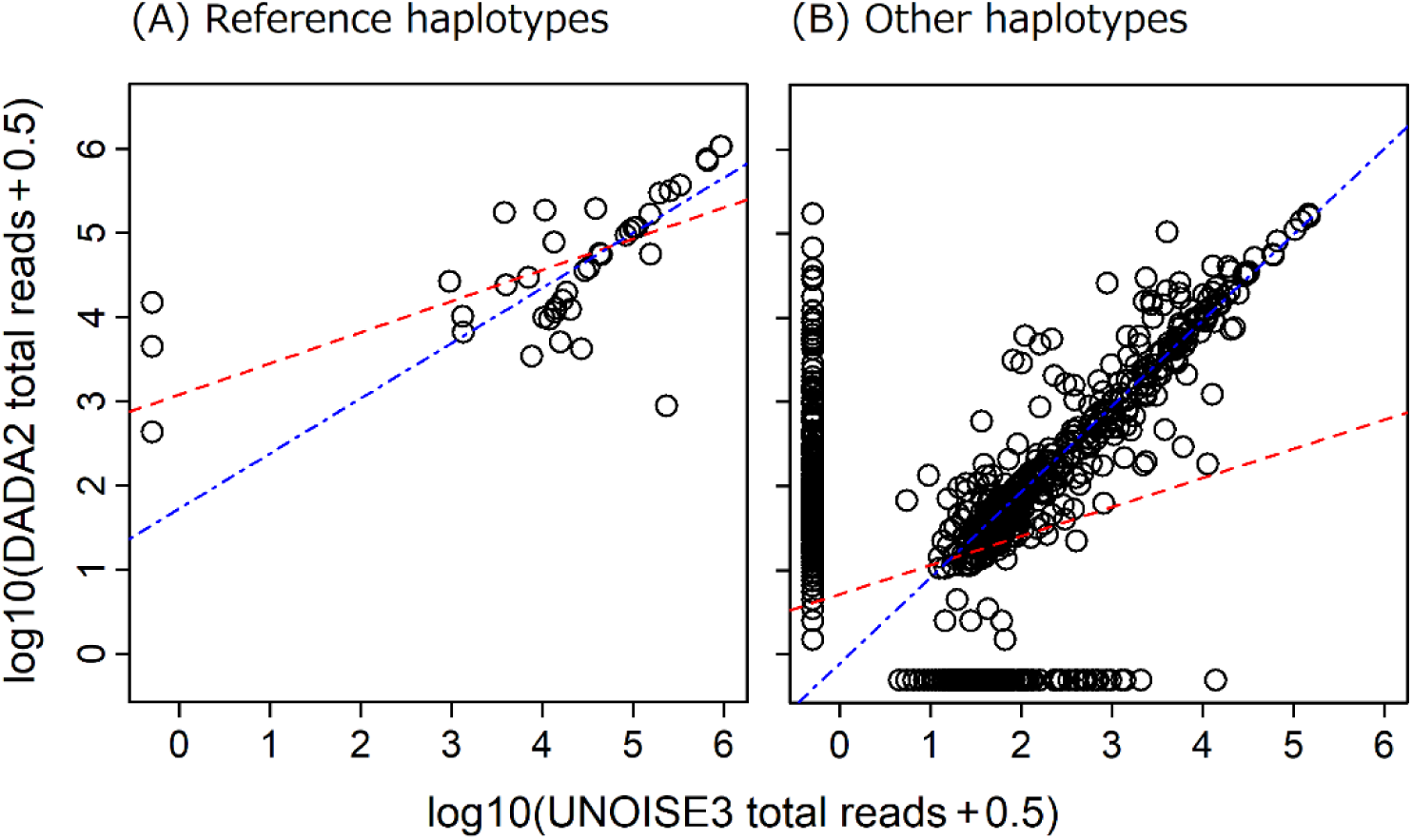
The relationship between total sequence reads of detected haplotypes by two denoising software packages, UNOISE3 and DADA2. (A) Reference haplotypes (total 41/42 haplotypes). (B) Haplotypes that did not correspond to any of the reference haplotypes. The 38/42 and 430/1,428 haplotypes were detected by UNOISE3 and DADA2, respectively. Red dashed and blue dot-dashed lines indicate the linear regression lines estimated using all detected haplotypes and only shared haplotypes detected by both UNOISE3 and DADA2, respectively.

### The haplotypes detected by DADA2

According to the GLMs, the detection probability of each haplotype among 20 filters or 15 PCR replications showed significant positive relationships with the number of owner specimens (*p* < 0.001, *p* < 0.001, respectively; Fig. 4). The increase in the number of detected reference haplotypes was gentle in both accumulation curves which were estimated using two data set including haplotypes obtained from 5 or 15 PCR replicates per filter (Fig. 5AB). Besides, the 95% confidence intervals of two accumulation curves overlapped considerably (Fig. 5AB). There was a significant positive relationship between the total read abundance of each haplotype and the number of specimens that owned the corresponding haplotype in the 96 captured Ayu specimens (lm, *p* < 0.01; Fig. 6).

**Fig. 4.**
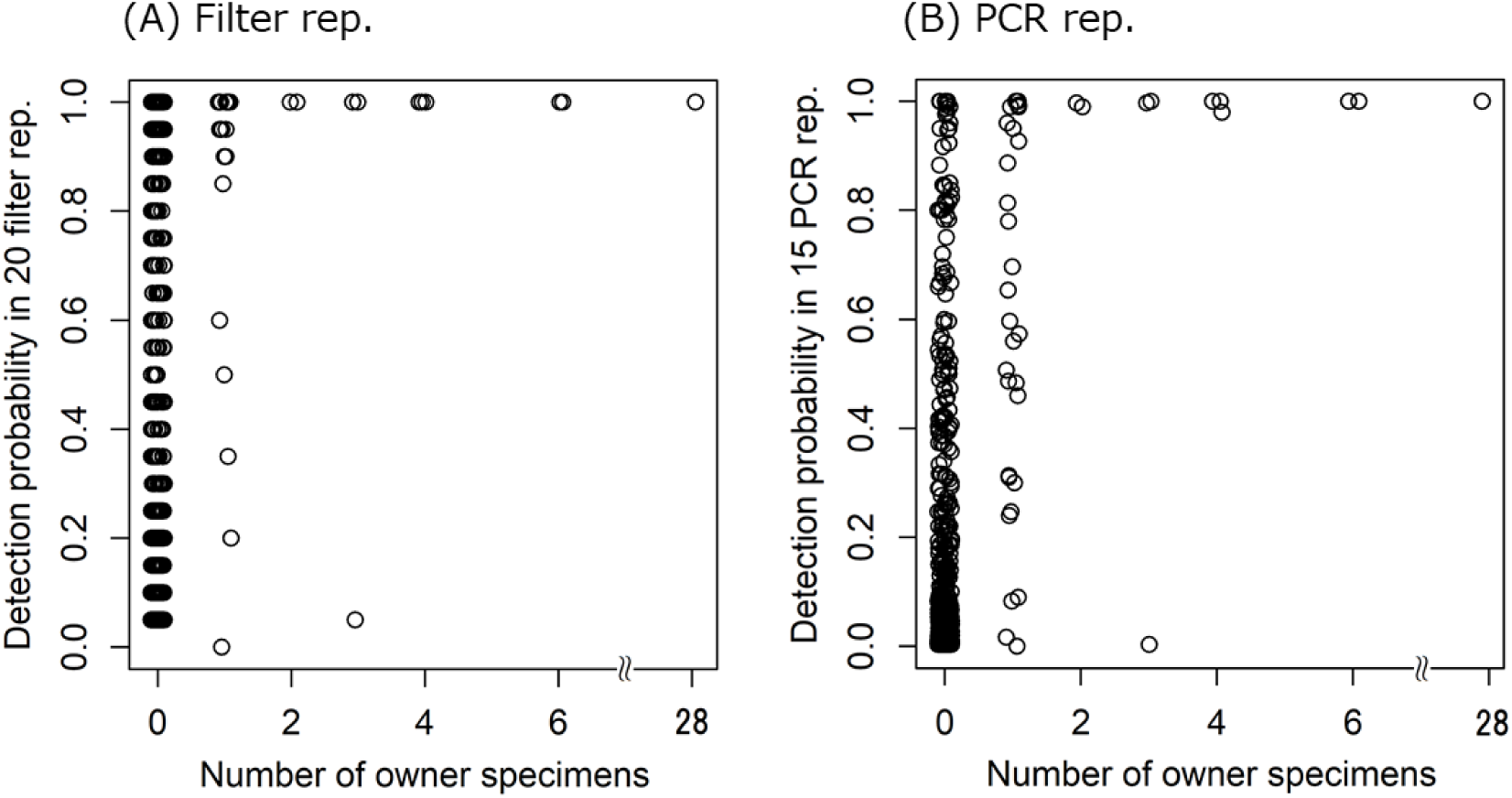
The relationship between the number of owner specimens and the detection probability: (A) in 20 filter replications, and (B) among 15 PCR replications. The haplotype detected from > 1 out of 15 PCR replications were defined as detected haplotypes from each filter. The detection probability of each haplotype on 20 filter or 15 PCR replications showed significant positive relationships with the number of owner specimens.

**Fig. 5.**
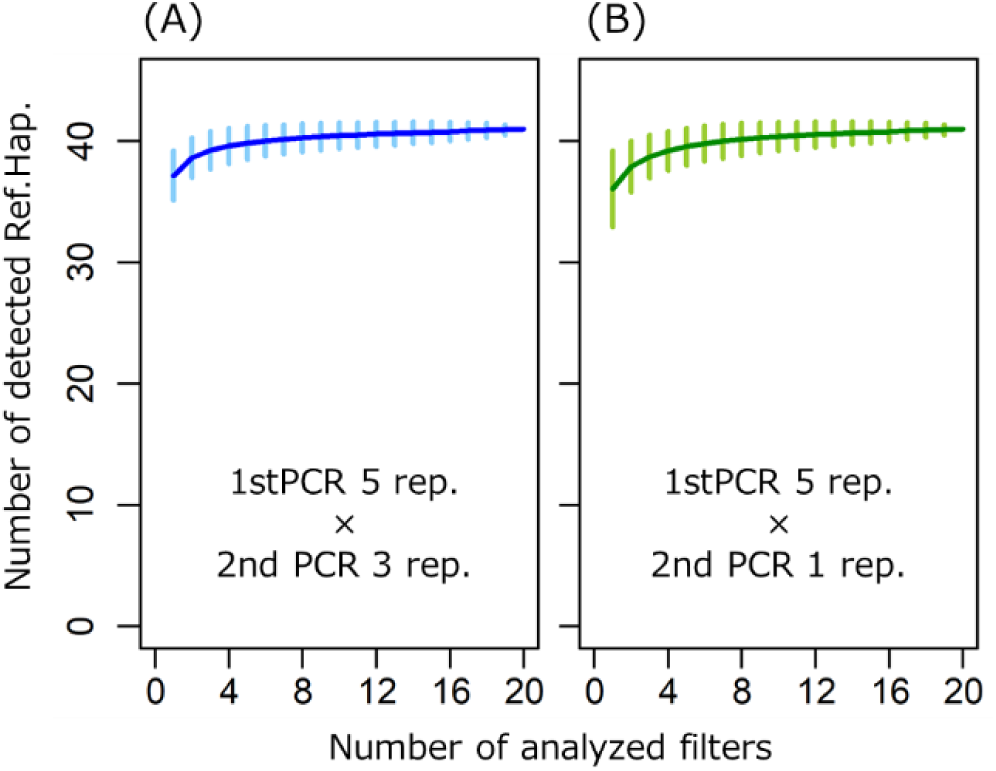
Accumulation curves of reference haplotypes estimated based on the number of analysed filters in eDNA analysis. (A) and (B) represent the accumulation curves estimated using the 15 and 5 PCR replications data (denoised by DADA2 denoising software package), respectively. Vertical bars indicate 95% confidence intervals.

**Fig. 6.**
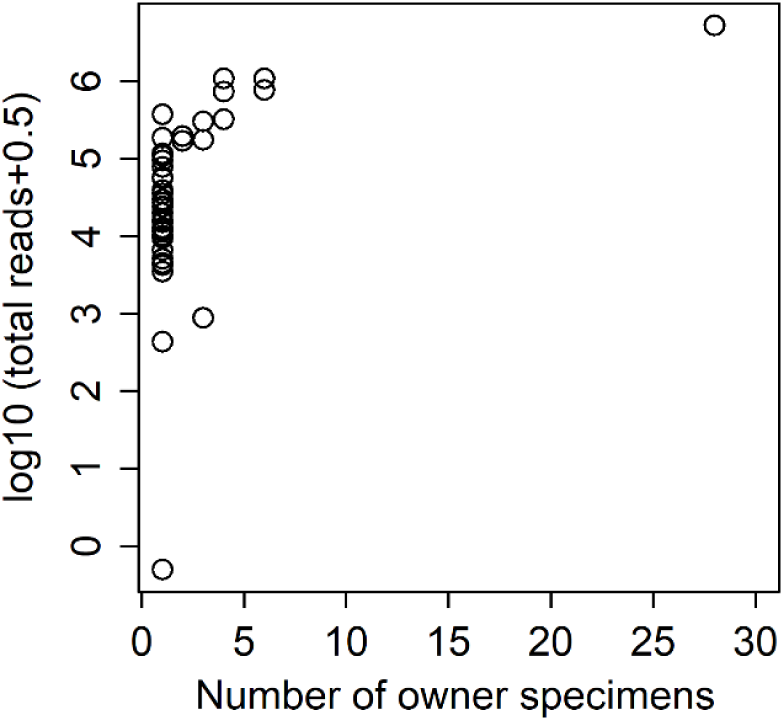
The relationship between the number of owner specimens and the total reads of each reference haplotypes.

## Discussion

Overall, we could obtain comparable results for the large-scale of capture-based conventional methods by using the eDNA-based method and demonstrated the usefulness of eDNA analysis for evaluating intraspecific genetic diversity in a wild fish population. The use of eDNA analysis enables us to significantly save time and labour for the survey. This advantage might promote the use of eDNA analysis for population genetics, phylogeography, and natural resource management. However, it is necessary to understand the risks of the false-positives and false-negatives for the accurate estimation of intraspecific diversity based on eDNA analysis in the field.

### Interpretation of intraspecific diversity data obtained with eDNA and conventional method

In eDNA analysis, 972 (UNOISE3) and 967 (DADA2) haplotypes were detected from 20 filter replications; however, this result should not be simply interpreted as eDNA analysis having greater detection efficiency than the capture-based conventional method, which was expected to detect about 250 haplotypes (Fig. 2B). In the previous study using tank water, some false positive haplotypes were presented in the final result after the denoising (Tsuji et al., 2019). Thus, some false positive haplotypes that were derived from erroneous sequences might still be included in the results, even if data here were denoised. Contrariwise, in the conventional method, the accumulation curve of detected haplotypes did not reach a plateau even when all captured specimens were analysed (Fig. 2A). When this study was performed, numerous individuals of Ayu were present at the sampling site because they were in active season for upstream migration from Lake Biwa (Azuma, 1973; Iguchi et al., 2002). Although it is highly likely that there was an underestimate because of the lack of specimens, we estimated that at least an additional 1,900 specimens are needed to evaluate the whole haplotype diversity at the sampling site (Fig. 2B). Therefore, it is likely that the results of the conventional method in this study (n = 96) did not cover the whole intraspecific genetic diversity of the Ayu population at the sampling site. Thus, eDNA analysis and the conventional method showed a tendency to overestimate and underestimate intraspecific genetic diversity, respectively.

### Comparison of detected haplotypes by eDNA analysis using UNOISE3 and DADA2

The DADA2 successfully detected more reference haplotypes than did UNOISE3 (Fig. 3A, Fig. S1), which suggests that the denoising of HTS data using DADA2 enables us to better estimate the presented haplotype at the sampling site. This difference in detection accuracy between UNOISE3 and DADA2 in this study is probably caused by the difference in the strategy of each denoising software. The UNOISE3 algorithm emphasized more on the sequence read abundance to denoise. eDNA of the rare haplotypes (i.e. where few individuals have the haplotypes) might exist at a low concentration in the water sample and they are more difficult to amplify by PCR. Thus, the numbers of sequence reads of the rare haplotypes were small and more likely to be eliminated. Although the sensitivity to the small difference is improved by increasing the alpha value, we do not recommend the major change of the alpha value because there is a trade-off between the sensitivity and the risk of increment of false positive haplotypes (Edgar 2016) (Table S2).

There was a significant positive relationship in the total reads of the detected unknown haplotypes between UNOISE3 and DADA2. Also, when only shared haplotypes of two denoising software packages were used, R^2^ values drastically improved (Fig. 3AB). When all of the reference haplotypes detected by UNOISE were also detected by DADA2, the haplotypes detected by both of these denoising software packages are more likely to represent haplotypes that are present in a water sample than unknown haplotypes detected by either of the two methods.

### Potential detection accuracy of eDNA analysis using DADA2

The eDNA-based method using DADA2 successfully detected 41 out of 42 reference haplotypes and demonstrated that eDNA analysis could evaluate intraspecific genetic diversity in the Ayu population, which were presented at the sampling time under field conditions. The detection probabilities of each reference haplotype among the 20 filters and 15 PCR replications increased with the number of owner specimens (Fig. 4A, B). Additionally, all but one reference haplotype owned by two or more specimens were detected from all filter replications with detection probability > 0.98 in 15 PCR replications. In contrast, reference haplotypes owned by one specimen were detected with various detection probabilities in filter and PCR replications. This result might be caused by the scarcity of eDNA molecules in the water sample when haplotypes were relatively rare in the population. However, filter replications have a smaller variation of detection probability than have PCR replications; 26 out of 30 reference haplotypes owned by one specimen were detected with probability > 0.8 among 20 filter replications. These results suggest that the haplotype selection based on the detection probability among multiple filter replications can eliminate haplotypes with a high probability of false positives. Furthermore, haplotypes detected detection probability of > 0.8 among 20 filter replications by only eDNA analysis (i.e. owner specimen was zero in the conventional method) might be derived from specimens that were not sampled by the conventional method. In this study, when only the haplotypes of with detection probabilities > 0.8 among 20 filter replications were selected, a total of 164 haplotypes including 36 reference haplotypes were detected by eDNA analysis. Considering that the estimated haplotype diversity based on the result of the conventional method was about 250 haplotypes, the 164 haplotypes that were detected by eDNA analysis with haplotype selection based on the detection probability appear to be a reasonable result.

### False-negative haplotype in eDNA analysis

When we performed denoising by DADA2, there was one false-negative result and some reference haplotypes with low detection probability in multiple filters and/or PCR replications (Fig. 4 and Fig. S1). It is considered that the false-negative result and decrease in detection probability among multiple replications of filters and PCRs were caused by three main reason: (1) failure of eDNA capture during the water sampling because of the scarcity or degradation of eDNA molecules (Evans et al., 2017), (2) failure of PCR amplification of eDNA in the sample because of scarcity of DNA molecules or the PCR inhibition (Jane et al., 2014, Ostberg et al., 2018), and (3) incorrect elimination in denoising of HTS data using denoising software based on the ASV method (Rosen et al., 2012).

To investigate the causes of the false-negative results in this study, all fastq files including raw reads were reanalysed without denoising. The base-calling errors were eliminated on the basis of the quality filtering, and the pre-processing and dereplicating of data was performed using a custom pipeline described by Sato et al. (2018). As a result, all reference haplotypes were obtained from the 20 filter replications; however, the detection probability of each reference haplotype among 15 PCR replications was varied from 33% to 100% (Fig. S1C). This result showed that eDNA molecules of all reference haplotypes would be included in all filter replications and amplified at least in some PCR replications. Additionally, sample water quality was equal for all filter replications; thus, the effect of PCR inhibition caused by water quality would be equal for all filter replications.

Furthermore, the low sequence read abundance that was caused by the failure of PCR amplification was likely to cause erroneous denoising by DADA2 (cf. Callahan et al., 2016). Therefore, in this study, the false negative result and the low detection probability on some reference haplotypes were considered to have been caused by the failure in PCR amplification and denoising of scarce eDNA.

In the reanalysis without denoising, much more haplotypes than the reference haplotypes (a total of 44,687) were detected from the 20 filter replications. Considering that 934 (UNOISE3) and 926 (DADA2) haplotypes other than reference haplotypes were detected when denoising was performed, it implies that the denoising software eliminated 97.90% and 97.93% of haplotypes, that have a high probability of false-positive results. Therefore, although there was some or one false negative haplotype, the denoising using UNOISE3 and DADA2 was considered indispensable for evaluating intraspecific genetic diversity by eDNA analysis.

### Future research suggestions

We found that the number of analysed filters and the number of PCR replications per filter have little effect on the number of detected reference haplotypes (Fig. 5AB). This finding suggests that many filters and PCR replications are not required in estimating intraspecific genetic diversity when using eDNA analysis. For example, when three filters that have five PCR replications are analysed, the detection efficiency of haplotypes will be equivalent to the situation where we analyse about 70 to 90 specimens by the capture-based conventional method (Fig. 2A, 5B). Although some false-negatives might occur in the eDNA-based method, those tend to be the rare haplotypes in the population. It is considered that the risk of false negatives in the eDNA-based method is lower than that in capture-based methods because 70 or more specimens are not generally analysed at one sampling site in capture-based methods. Therefore, we recommend the use of the eDNA-based method as an alternative or screening method for capture-based methods in large-scale surveys (e.g. in a wide area and/or in many sampling points) of intraspecific genetic diversity. An eDNA-based method would efficiently provide valuable information for estimating and evaluating intraspecific genetic diversity in a population.

We found a significant positive relationship between the total reads of each haplotype and the number of owner specimens, consistent with the results in previous studies using tank water (Tsuji et al., 2019) and field water (Sigsgaard et al., 2016). Although there are some issues regarding the use of sequence read abundance as an index of the abundance/biomass of target species (Ushio et al., 2018), the read abundance might reflect the quantitative relationships among haplotypes in a population. In general, the number of sequence reads obtained by HTS does not necessarily reflect the number of DNA copies. However, it was recently shown that the inclusion of internal standard DNA in the eDNA sample enables the simultaneous determination of the quantity and identity of the eDNAs of multiple species (Ushio et al., 2018). Thus, the application of the quantitative technique for HTS technology could enable the quantitative evaluation of intraspecific diversity by eDNA analysis in future studies. Besides, false positive haplotypes could be eliminated by the cutting off haplotypes with < 1 copy per sample volume used for PCR, and the degree of overestimation of intraspecific genetic diversity by the eDNA-based method may be suppressed.

The eDNA-based evaluation of intraspecific genetic diversity will be used more widely for the survey of intraspecific genetic diversity, population genetics and phylogeography if the technique becomes more developed and refined in the future. This study showed that the eDNA-based method detected various haplotypes comparable with those observed in large-scale capture surveys, and can evaluate intraspecific genetic diversity in a wild fish population from the field sample.

## Acknowledgements

We thank Mr. Sakurai, S., Mr. Kakimi, N., Mr. Hongo, M., Mr. Sogo, Y., and Mr. Motozawa, H. (Ryukoku University) for supporting the field survey. This work was supported by JSPS KAKENHI Grant Number 16K18610 and Grant-in-Aid for JSPS Research Fellow Grant Number JP18J10088.

## Author contributions

i. the conception or design of the study: S.T., and H.Y
ii. the acquisition, analysis, or interpretation of the data: S.T., A.M., M.M., M.U., H.S., T.M., and H.Y.
iii. writing of the manuscript: S.T., A.M., M.M., M.U., H.S., T.M., and H.Y.

## Data accessibility statement

The minimal raw dataset is uploaded to the DDBJ Sequence Read Archive (https://www.ddbj.nig.ac.jp/dra/index-e.html; Accession number: DRA009149)

## Supporting Information

**Fig. S1.**
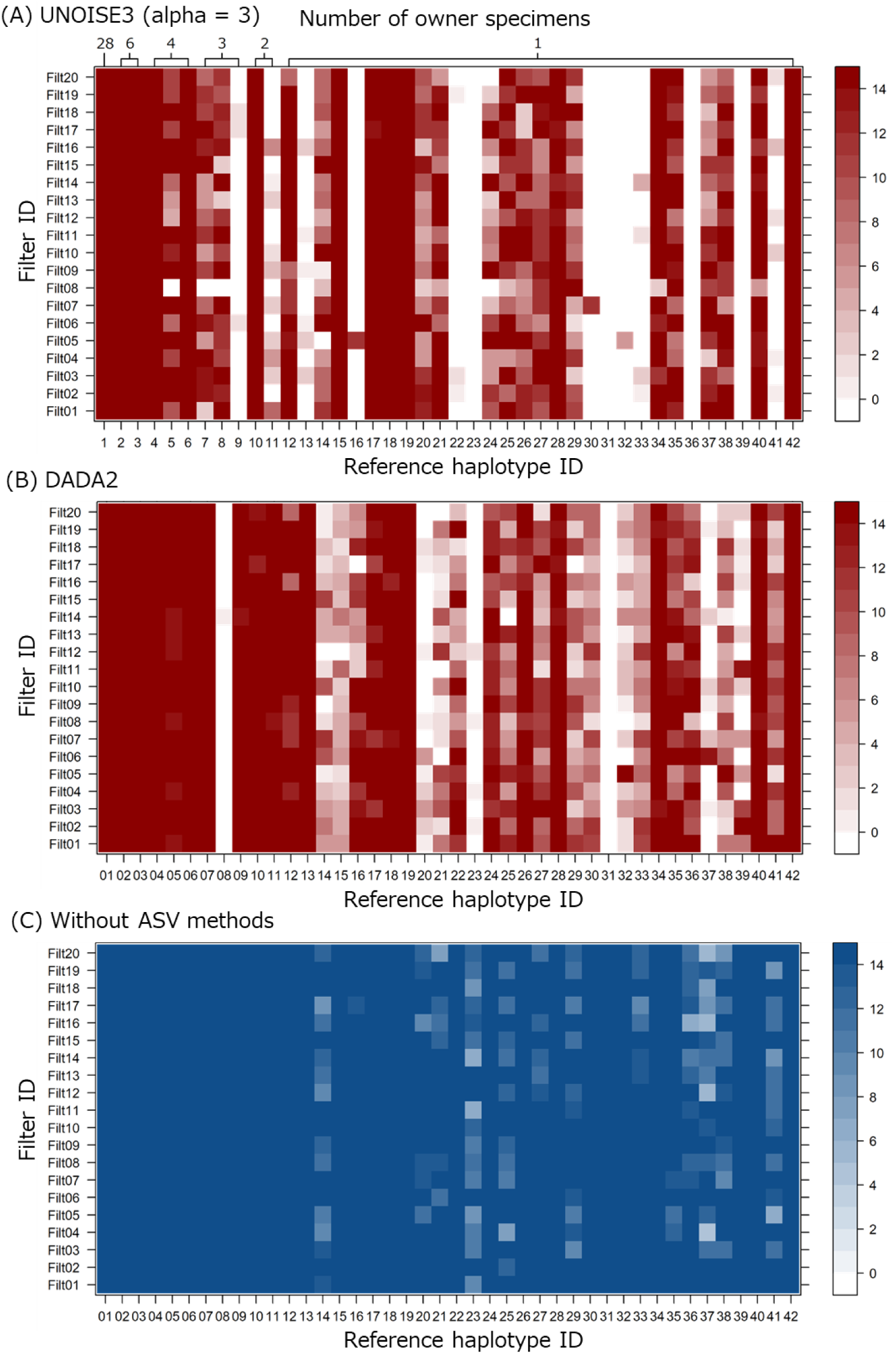
Level maps depicting the number of detections among 15 PCR replications of each reference haplotype per filter. Each of level map indicates the result obtained when denoising was performed by (A) UNOISE3, (B) DADA2, and (C) not performed (analysed using a custom pipeline, Sato et al. 2018), respectively. Horizontal axes indicate the reference haplotype ID and number of owner specimens of each reference haplotype, respectively.

